# Predictive encoding of motion begins in the primate retina

**DOI:** 10.1101/2020.09.10.291419

**Authors:** Belle Liu, Arthur Hong, Fred Rieke, Michael B. Manookin

**Affiliations:** Department of Physiology and Biophysics, University of Washington, Seattle, WA 98195; Department of Physics, National Tsing Hua University, Hsinchu, Taiwan; Interdisciplinary Program of Science, National Tsing Hua University, Hsinchu, Taiwan; Department of Ophthalmology, University of Washington, Seattle, WA 98109; Vision Science Center, University of Washington, Seattle, WA 98109

**Keywords:** neural coding, motion prediction, spatiotemporal correlations, retina

## Abstract

Survival in the natural environment often relies on an animal’s ability to quickly and accurately predict the trajectories of moving objects. Motion prediction is primarily understood in the context of translational motion, but the environment contains other types of behaviorally salient motion, such as that produced by approaching or receding objects. However, the neural mechanisms that detect and predictively encode these motion types remain unclear. Here, we address these questions in the macaque monkey retina. We report that four of the parallel output pathways in the primate retina encode predictive information about the future trajectory of moving objects. Predictive encoding occurs both for translational motion and for higher-order motion patterns found in natural vision. Further, predictive encoding of these motion types is nearly optimal with transmitted information approaching the theoretical limit imposed by the stimulus itself. These findings argue that natural selection has emphasized encoding of information that is relevant for anticipating future properties of the environment.

## INTRODUCTION

Extracting motion information from the environment is a key function of the visual system. Motion produces strongly stereotyped patterns of light stimulation on the retina, and these patterns (correlations) comprise a rich source of behaviorally salient information. Pairwise correlations are formed between two points in space and time during translational motion (Hassenstein and Reichardt, 1956; Adelson and Bergen, 1985) and accessing these correlations allows some retinal circuits to anticipate the future position of moving objects, a function which we refer to as predictive motion encoding (Palmer et al., 2015; Berry et al., 1999; Schwartz et al., 2007; Johnston and Lagnado, 2015; Leonardo and Meister, 2013). However, the natural environment contains strong asymmetries between the distribution of dark and bright light intensities and these asymmetries produce correlations between three or more points in space and time during natural motion, such as when an object becomes closer (diverging correlations) or farther away (converging correlations) (Field, 1987; Dong and Atick, 1995). These higher-order correlations constitute a potentially rich source of information for estimating visual motion (Clark et al., 2014; Fitzgerald et al., 2011; Sayrol et al., 1996; Anderson and Giannakis, 1995; Nitzany and Victor, 2014; Nitzany et al., 2016).

Retinal mechanisms that estimate future motion are best understood in the context of deterministic motion in which an object translates smoothly across the retina (Berry et al., 1999; Schwartz et al., 2007; Johnston and Lagnado, 2015; Leonardo and Meister, 2013). Under these conditions, estimating the future position of a moving object is relatively straightforward. However during natural vision, object motion generally includes strongly random elements. For example, a viewer’s own motion from head and eye movements can produce large and frequent displacements of visual input on the retina (Kuang et al., 2012; Van Der Linde et al., 2009), and these fluctuations pose a considerable computational challenge for neural circuits tasked with estimating the future trajectory of a moving object (Palmer et al., 2015). Thus, understanding motion processing in natural environments requires knowledge of how neural circuits encode pairwise and higher-order correlations in the presence of random fluctuations.

To address these questions, we measured the responses of several neural types in the macaque monkey retina to stochastic stimuli containing distinct classes of motion correlation (Hu and Victor, 2010; Nitzany and Victor, 2014). We report that parasol and smooth monostratified ganglion cells encode predictive information about both pairwise and higher-order motion correlations in their outputs. The encoding of this information about future motion is nearly ideal—neural responses represent virtually all of the predictive information available in the stimuli. Further experiments reveal that this predictive encoding is present at the synaptic output of diffuse bipolar cells and emerges downstream of horizontal cells in the outer retina. Thus, the primate visual system begins encoding of information that can be utilized to predict future motion at the second visual synapse.

## RESULTS

### Primate ganglion cells encode higher-order motion correlations

Natural environments contain asymmetrical distributions of dark and bright light intensities and this asymmetry produces higher-order spatiotemporal correlations that are useful for estimating visual motion (Fitzgerald et al., 2011). Indeed, humans use these correlations in motion estimation and this sensitivity is thought to arise in the visual cortex (Hu and Victor, 2010; Clark et al., 2014). To determine whether the primate retina encoded these motion patterns, we measured the responses of ganglion cells, the retinal output neurons, to several classes of motion stimuli. Neural responses were measured to stimuli containing correlations between two points or three points in space and time (Hu and Victor, 2010).

Two general classes of three-point correlations were tested: 1) diverging correlations that consisted of a point in space that diverged to two points in space at a later time, and 2) converging correlations where two points in space converged to a single point at a later time (Figure 1). These types of correlations can arise when objects approach (diverging correlations) or recede away from an observer (converging correlations) (Fitzgerald et al., 2011; Clark et al., 2014; Hu and Victor, 2010; Leonhardt et al., 2016; Nitzany and Victor, 2014; Appleby and Manookin, 2020). These three-point correlations were further subdivided into positive or negative based on the contrast polarity of the diverging or converging points. In all cases, the three-point motion stimuli lacked two-point spatiotemporal correlations (Figure 1B) (Hu and Victor, 2010; Clark et al., 2014).

**Figure 1.**
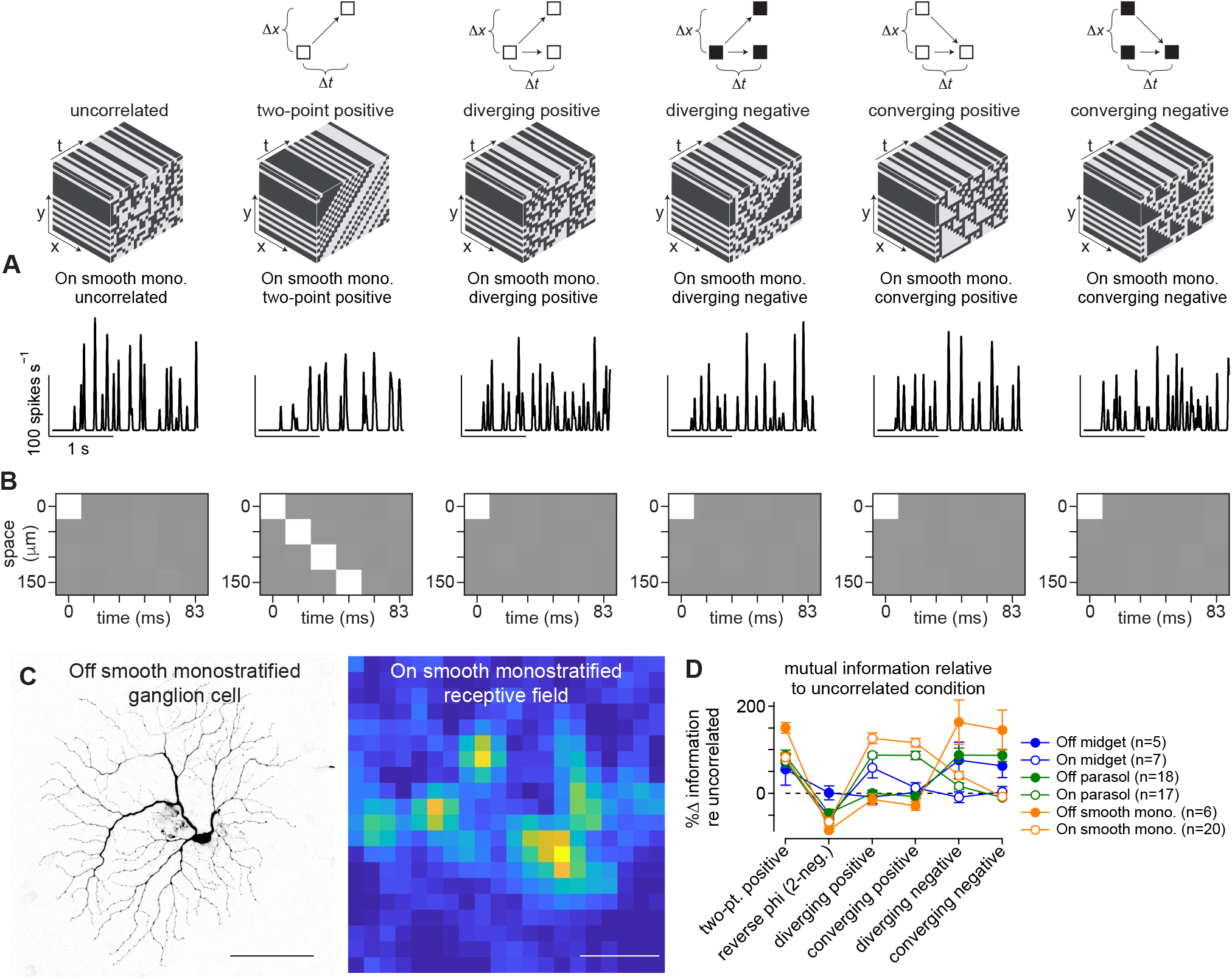
Ganglion cells show sensitivity to higher-order correlations. (A) *Top*, Example space-time plots for six of the stimulus classes used in the study. *Bottom*, Spike rate (in spikes s^−1^) as a function of the time for these stimuli in an On smooth monostratified ganglion cell. (B) Autocorrelation for five of the stimulus types showing space (*y*-axis) as a function of time (*x*-axis). (C) Morphology from an Off-type (*left*) and receptive field from an On-type (*right*) smooth monostratified ganglion cells recorded as part of this study. The receptive field showed characteristic hotspots. Scale bar, 0.1 mm. (D) Peak values for the time-shifted mutual information for six types of motion correlations relative to the information for the uncorrelated stimulus. Data are shown for several ganglion cell types. Circles and error bars indicate mean ± SEM.

We compared responses to these motion stimuli to a stimulus lacking spatiotemporal correlations (uncorrelated condition; Figure 1). We estimated the amount of information that the response in one interval of time encoded about the stimulus in another interval of time—the time-shifted mutual information (bin width, 16.7 ms; see Methods; Figure S1). This method measures the amount of information that can be obtained about a stimulus from neural responses— a stimulus that produces increased information rates relative to the uncorrelated condition indicates that neural responses encode more information about that stimulus. Further, because the three-point motion stimuli vary from the uncorrelated stimulus only in their higher-order correlations, we interpret greater information rates for these stimuli as evidence that a cell encodes higher-order correlations (Figure 1B). We compared the peak values for the time-shifted mutual information across stimulus conditions and cell types.

Smooth monostratified and parasol ganglion cells showed peak information rates that were ∼2-fold (100%) higher relative to the uncorrelated stimulus for three-point motion stimuli aligned with the contrast polarity of the cell—positive contrasts for On cells and negative contrasts for Off cells (Figure 1E; p ≤ 0.02; Wilcoxon signed rank test). These increased information rates for the three-point motion correlations indicate that smooth monostratified and parasol cells encode these correlations in their spike outputs. We next investigated the dependence of this encoding on spike timing.

### Information encoding occurs with high temporal precision

The analysis presented in Figure 1 calculated the time-shifted mutual information for a time window corresponding to the stimulus update rate (Δ*t*, 16.7 ms). However, many neurons can encode stimulus features on timescales of only a few milliseconds (Reinagel and Reid, 2000; Froemke and Dan, 2002; Gollisch and Meister, 2008; Uzzell and Chichilnisky, 2004; Bialek et al., 1991; Rieke et al., 1997; de Ruyter van Steveninck et al., 1997; Panzeri et al., 2001). To determine the relationship between spike timing and the encoding of motion correlations, we calculated the mutual information between the stimulus and spike response of a cell after randomly varying spike times. Spikes were binned in 1-ms intervals and the timing of each spike was determined by drawing a random value from a Gaussian distribution with a mean centered at the time of the original spike and a standard deviation between 0–100 ms (see Methods). This process was repeated 50 times for each standard deviation (Figure 2).

**Figure 2.**
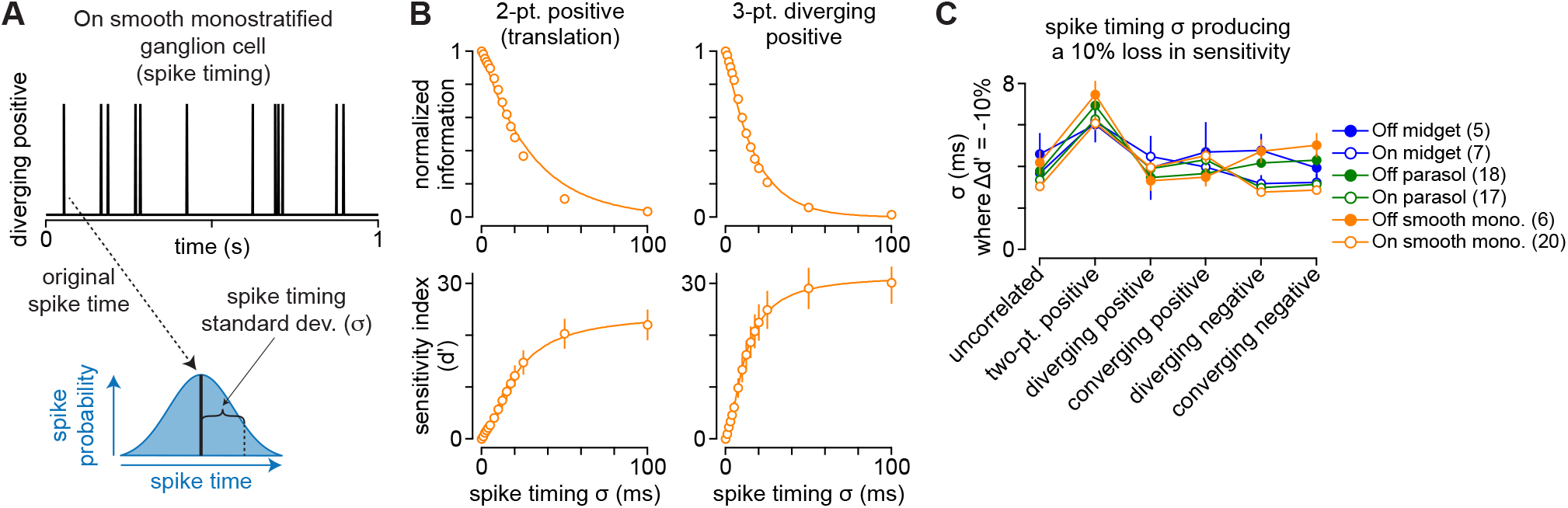
Information encoding occurs on short time scales. (A) Example spike train for a one-second portion of a stimulus containing diverging positive correlations (*top*). To measure the relationship between spike timing and information transmission, each spike was randomly shifted in time. Time shifts were drawn from a Gaussian distribution with a mean corresponding to the timing of the original spike and a standard deviation (*σ*, *bottom*). The process was repeated for each spike train in each cell at multiple standard deviations (0-100 ms). (B) Normalized information (*top*) and sensitivity index (*d′*, *bottom*) as a function of the standard deviation of spike timing (*x*-axis) across On smooth monostratified ganglion cells (n = 20). Solid lines show either a single-exponential fit (*top*) or a sigmoidal (*bottom*) to the data. (C) Spike timing standard deviation at which the sensitivity index increased by 10% relative to the condition with spike time unaltered (*σ*, 0 ms). Data are shown across six stimulus conditions and six ganglion cell types. Circles and error bars indicate mean ± SEM.

Spike timing could affect information across a range of time scales. If information encoding were determined solely by the statistics of the stimulus, information should decrease only when the variation in spike timing exceeded the update interval of the stimulus itself (16.7 ms). However, if encoding occurred on finer timescales, information should decrease rapidly with fine-grained changes in spike timing. Our analysis supported the latter premise that these primate ganglion cell types encoded information on relatively fine time scales. Encoded information decayed rapidly after shifting spike timing by only a few milliseconds (Figure 2B, *top*). This decline continued such that, at time shifts of 50 ms, information about the stimulus was almost completely degraded.

To determine the threshold at which variation in spike timing significantly degraded information, we calculated the sensitivity index (*d′*) between the original and shifted spike trains (see Methods). Lower *d′* values indicate that the information content of the shifted condition was similar to the original spike train, whereas larger index values indicate that information between the two spike trains is more distinct (i.e., degraded). The sensitivity index increased rapidly at time shifts between 1–20 milliseconds, consistent with a rapid degradation in information. We defined a significant change in information as a 10% increase in *d′* relative to the saturating level. Across cells and stimulus conditions, this threshold was typically reached in less than five milliseconds, indicating that encoding these stimuli requires a high level of temporal precision (Figure 2C).

### Time scales of encoding match correlation structure of stimuli

The correlations inherent to visual motion mean that the motion trajectory leading up to a particular time (*t* ≤ 0) provides information about the likely trajectory of motion at later time points (*t >* 0). This information can then be used to estimate motion trajectories and to guide behavior (Bialek et al., 2006; Palmer et al., 2015; Salisbury and Palmer, 2016; Chalk et al., 2018; Sederberg et al., 2018). The correlations in the motion stimuli used in our study contained this type of predictive information within a time window of approximately ±50 ms (Figure 3D). To determine whether the timescales of encoding match the time course of available information in the stimulus, we calculated the mutual information between a cell’s spike output and the stimulus at different time lags relative to the stimulus presentation (Figure S1). We then compared these results to the measured mutual information between the stimulus and itself at the same time lags. The temporal profile of information available in the stimulus was shifted to account for the time lag between the presentation of the stimulus at the level of the photoreceptors and the ganglion cell output (∼30–50 ms; Figure 3A). This time lag was calculated directly from the spatiotemporal receptive field measured for each cell (see Methods).

**Figure 3.**
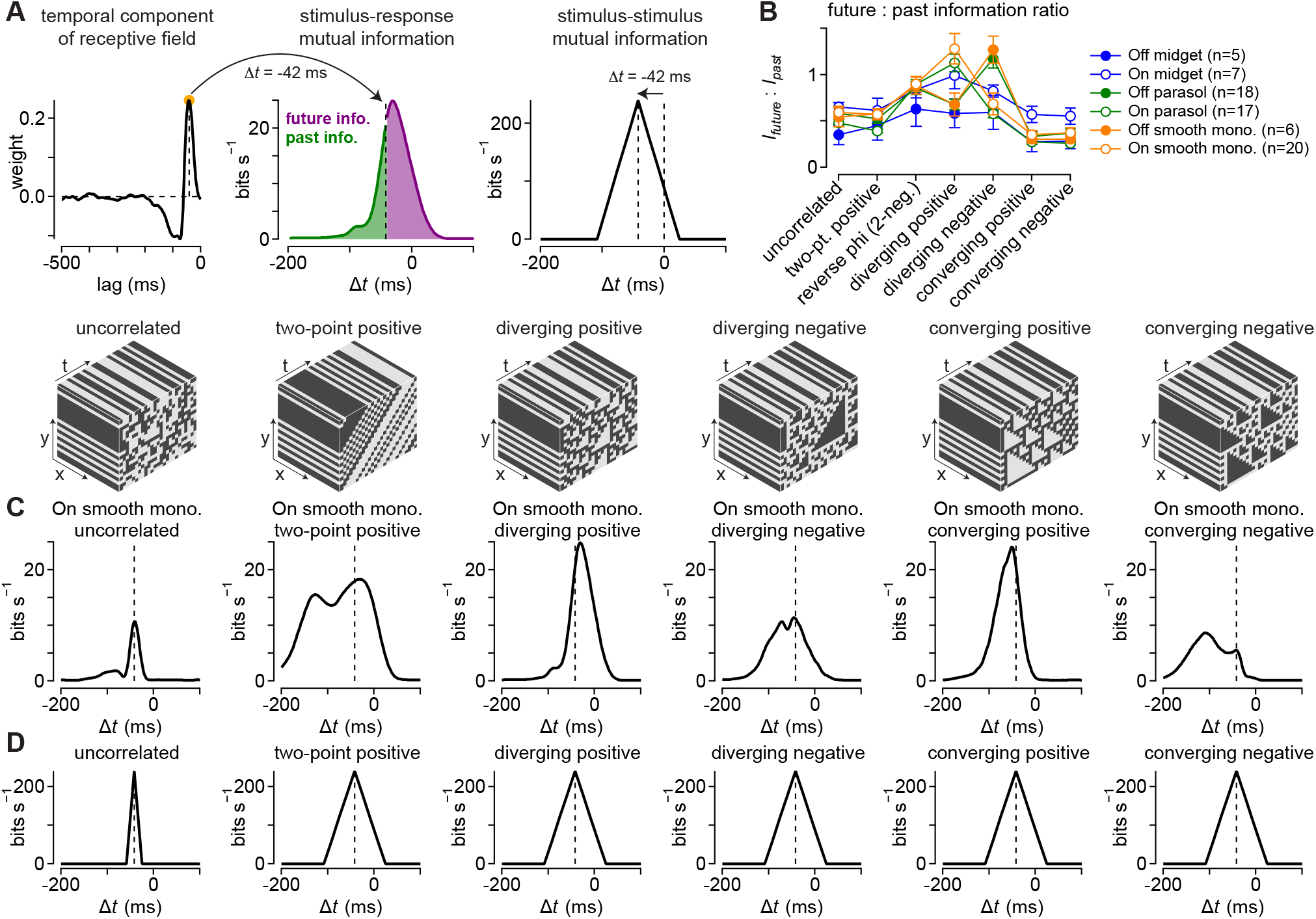
Time scales of encoding match higher-order correlations. (A) The spatiotemporal receptive field was measured for the uncorrelated stimuli in each cell. The peak (On cells) or trough (Off cells) of the temporal receptive field was used as the measure of lag in visual processing and was defined as *t* = 0 for subsequent measures of past and future information. (B) Ratio of future to past information for six primate ganglion cell types across the seven stimulus classes used in this study. (C) Information rate (in bits s^−1^) as a function of the time delay (in ms) for six stimulus classes in an On smooth monostratified ganglion cell. (D) Information that each stimulus class contains about itself as a function of the time delay. The peak was shifted relative to the measured lag for signal transmission from the photoreceptors to the ganglion cell output.

This analysis for an On smooth monostratified ganglion cell is shown in Figure 3. The cell’s encoding of the stimulus closely matched the time course of the information contained within the stimulus itself. For the uncorrelated noise condition, which contained information about itself only within the window of a single frame (∼17 ms), the cell encoded stimulus information in a relatively short time window (Figure 3C, *far left*). These time windows became broader for encoding of motion stimuli, indicating that the time course over which the cell encoded these stimuli roughly matched the time course of available information contained within the motion stimuli (Figure 3C, D).

In addition to tracking the correlation structure in the stimuli, information encoded in the cell’s responses was shifted in time relative to the stimulus presentation for pairwise and diverging motion stimuli (Figure 3B, C). Encoding of diverging correlations and two-point positive correlations was shifted to future (i.e., positive) time lags. This shift indicates that the cell’s response at a given time reliably predicts the pattern of motion at later time points. This pattern in which a cell’s responses anticipate subsequent motion is characteristic of predictive motion encoding (Palmer et al., 2015; Berry et al., 1999; Johnston and Lagnado, 2015).

The observed displacement in the time-shifted mutual information towards future time lags could arise from the effects of motion correlations on the retinal circuit, but could also arise from other aspects of the stimulus. To distinguish between these possibilities, we extracted the motion correlations from the stimulus and used these correlations in calculating the time-shifted mutual information (Figure S2). If the shift in mutual information were caused by a component of the stimulus other than the motion correlations, then the shift towards future time lags would be eliminated if the analysis were done on these correlations. Motion correlations were calculated from the products of the intensity values at the space and time intervals for each stimulus class (Nitzany and Victor, 2014). For example, pairwise motion correlations were determined by multiplying the stimulus intensity at one point in space with the intensity at an adjacent point in space on the next stimulus frame (Hassenstein and Reichardt, 1956). After computing the local motion correlations, we computed the time-shifted mutual information between these values and the cell’s spike output.

Indeed, this analysis showed the same pattern of predictive motion encoding as we observed in Figure 3. These results confirm that the predictive displacement in the time-shifted mutual information for pairwise and diverging correlations arises from the motion correlations in the stimulus. However, this predictive encoding did not occur for all motion correlations tested. The predictive shift was not present for converging correlations, as encoded information was not shifted to positive time lags for these stimuli (Figure 3). This finding is consistent with previous work demonstrating that different correlation classes differentially shape the response properties of parasol and smooth monostratified ganglion cells (Manookin et al., 2018; Appleby and Manookin, 2020). In the next section, we further investigate this relationship between stimulus classes by determining how closely cellular encoding of predictive information in these stimuli approximates an optimal encoding.

### Predictive encoding nearly optimal for pairwise and diverging correlations

A single neuron cannot encode all of the information available in a sensory input. For example, the uncorrelated stimulus used in the present work contained ≥ 2^15^ possible patterns within a given stimulus interval (∼16.7 ms). However, in the same time window, the ganglion cells in this study represented information with only between 0–7 spikes—2^3^ response levels. This means that only a small fraction of the information available in the stimulus was encoded in the spike output of these cells. Thus, the encoding scheme adopted by a neuron creates a compressed representation of the incoming stimulus with most information discarded (i.e., lossy compression; Figure 4A). Further, based on this compression scheme, one can determine the maximal amount of predictive information that can be encoded given the amount of information encoded about the past stimulus trajectory (Tishby et al., 1999). The predictive information in neural responses can then be compared to this optimal encoding (Figure 4B).

**Figure 4.**
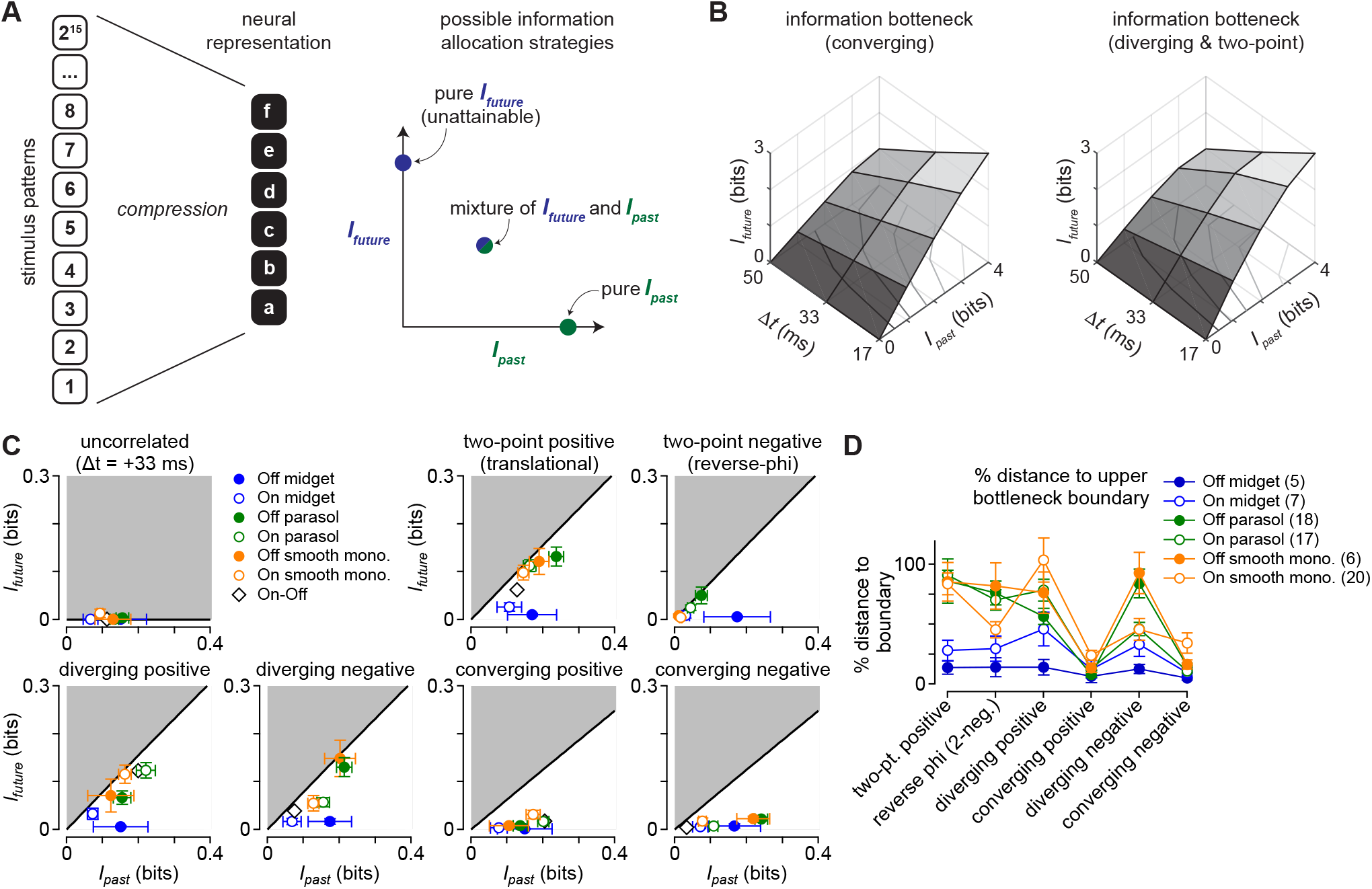
Encoding of predictive information is nearly optimal for certain stimulus classes. (A) *Left*, A potential lossy information encoding scheme that could be employed to represent components of a high-dimensional stimulus in the neural output. *Right*, This neural representation can allocate bandwidth for encoding information about the past stimulus history (*I_past_*) and information that can be used to anticipate future motion (*I_future_*). (B) Information bottleneck boundary for available past and future information at three future time lags. Curves are shown for the three-point converging stimuli (*left*) and the three-point diverging and two-point stimuli (*right*). (C) Encoding of predictive information (*I_future_*; *y*-axis) versus past information (*I_past_*; *x*-axis) for the seven stimulus classes. The black solid line indicates the theoretical boundary for predictive information and the shaded region defines the theoretically forbidden values. Predictive information was nearly optimal for two-point positive, diverging positive, and diverging negative correlations. Cellular representations of converging correlations lacked similar predictive information. (D) Encoded future information (*I_future_*) in six cell types relative to the theoretical maximum value defined by the information bottleneck. Values are given as a percentage of the boundary value for the six stimulus classes used in this study. Circles and error bars indicate mean ± SEM.

The results of this analysis are shown in Figure 4— the solid black line delineates the boundary for optimal predictive encoding (*I_future_*) as a function of the information about the past stimulus trajectory (*I_past_*). The shaded regions to the upper left define values for predictive information that are not theoretically attainable. Indeed, the entire predictive domain of the curve was unattainable for the uncorrelated noise stimulus, which lacked predictive information, and this was also reflected in a lack of measured predictive information (Figure 4C, *upper left*).

For the motion stimuli, which contained predictive information, the relationship between cellular encoding of this information and the optimal boundary varied both with cell type and stimulus type. As suggested by the data in Figure 3, ganglion cells encoded predictive information for two-point correlations and three-point diverging correlations. Further, this encoding was nearly ideal for On and Off smooth monostratified and parasol ganglion cells, which captured between 70–100% of the available predictive information for a contrast-aligned stimulus. This indicates that the spike output of these cells retains nearly all of the predictive information contained in these stimuli. Midget (parvocellular-projecting) ganglion cells, however, encoded relatively less predictive information in their spike outputs to these stimuli. In these cells, mean values fell between 38-80% for the two-point and diverging correlations, and predictive encoding was particularly low for Off midget cells. Thus, predictive encoding varied with ganglion cell type.

Unlike the two-point and diverging motion correlations, cellular encodings of converging motion correlations fell far off the optimal boundary (Figure 4C). Parasol and smooth monostratified cells encoded less than 50% of the available predictive information for these stimuli. This lack of strong predictive encoding for converging correlations is consistent with our previous work demonstrating a preference for approaching motion in these cell types. Approaching motion, which produces diverging correlations on the retina, elicited significantly larger responses than receding motion, which produces converging correlations (Appleby and Manookin, 2020). The data presented in the current study go further to indicate these neurons encode predictive information about diverging correlations that could be utilized to anticipate future motion trajectories.

### Predictive encoding nearly optimal at low contrast

Due to noise in the photoreceptors, decreasing the contrast of a stimulus also diminishes the signal-to-noise ratio. Such decreases in signal-to-noise ratio can severely limit the accuracy of information encoding (Abbott and Dayan, 1999). To determine whether encoding of predictive information was diminished at low contrast, we calculated past and future information encoded in the same cell at three different stimulus contrasts (contrasts: 0.25, 0.5, 1.0). Measurements were performed for uncorrelated, two-point motion, and three-point diverging motion stimuli in the same cell. We found that the encoded past and future information varied with stimulus contrast (Figure 5). Information calculated for time lags of Δ*t* = 0 (*I_past_*), 17, 33, and 50 ms showed differences in relative saturation at high contrast. Past information gradually increased with contrast and showed only weak saturation at the highest contrast level (Figure 5B). Saturation was greater for encoded future information and this trend increased with increasing time in the future such that, for the longest lag (+50 ms), the information encoded at the lowest and highest contrast levels depended only weakly on contrast. This saturation in future information resulted in a higher ratio of future to past information encoding at lower contrasts— a trend that again was strongest for time lags farther into the future (Figure 5C).

**Figure 5.**
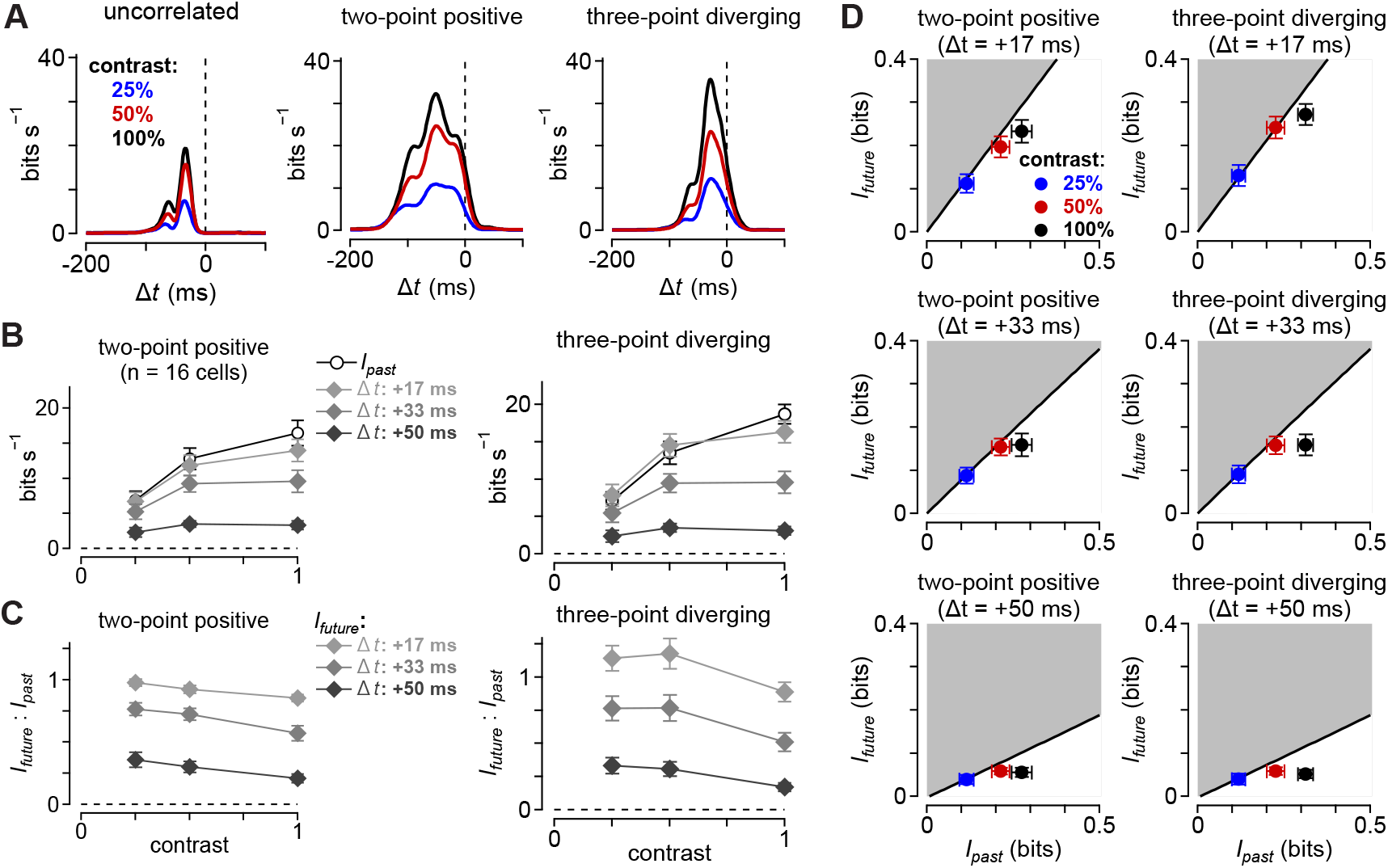
Ganglion cells optimally encode future information at low contrast. (A) Information rate (in bits s^−1^) as a function of the time delay (in ms) for three stimulus classes in an On smooth monostratified ganglion cell. Results are shown at three different contrasts for each stimulus class. (B) Relationship between encoded information as a function of stimulus contrast. Results are shown for past information and three future time lags. (C) Ratio of future (*I_future_*) to past (*I_past_*) information as a function of contrast in the same cells as in (B). Encoding of predictive information (*I_future_*; *y*-axis) versus past information (*I_past_*; *x*-axis) in 16 parasol and smooth monostratified cells. Measurements were taken at three different stimulus contrasts and three time lags. Circles and error bars indicate mean ± SEM.

We next asked whether the greater saturation in future information affected the relationship between cellular encoding and the optimal boundary defined by the information bottleneck. We found that this relationship varied with stimulus contrast and time lag. At a time lag of +17 ms, encoding of future information was near the optimal boundary at all contrasts (Figure 5D, *top*). At later time lags, however, the lowest contrast stimulus maintained a position near the optimal boundary, but the higher contrasts separated from this boundary. These data indicate that, at low contrast, parasol and smooth monostratified cells encode nearly all of the available future information for pairwise and diverging motion. Further, these cells most optimally encoded predictive motion information at low contrast, when the signal-to-noise ratio was the lowest. The implications of these results are addressed in the Discussion. We next investigated the circuit mechanisms mediating the encoding of predictive information.

### Predictive encoding of motion originates early in the visual pathway

The data presented thus far indicate that parasol and smooth monostratified ganglion cells encode predictive information about visual motion in their spike outputs. We next investigated the mechanisms mediating this predictive encoding. To do this, we assayed three different points in the retinal circuit: 1) the voltage responses of horizontal cells in the outer retina, 2) amacrine cell inhibitory synaptic outputs, and 3) diffuse bipolar cell excitatory synaptic outputs (Figure 6).

**Figure 6.**
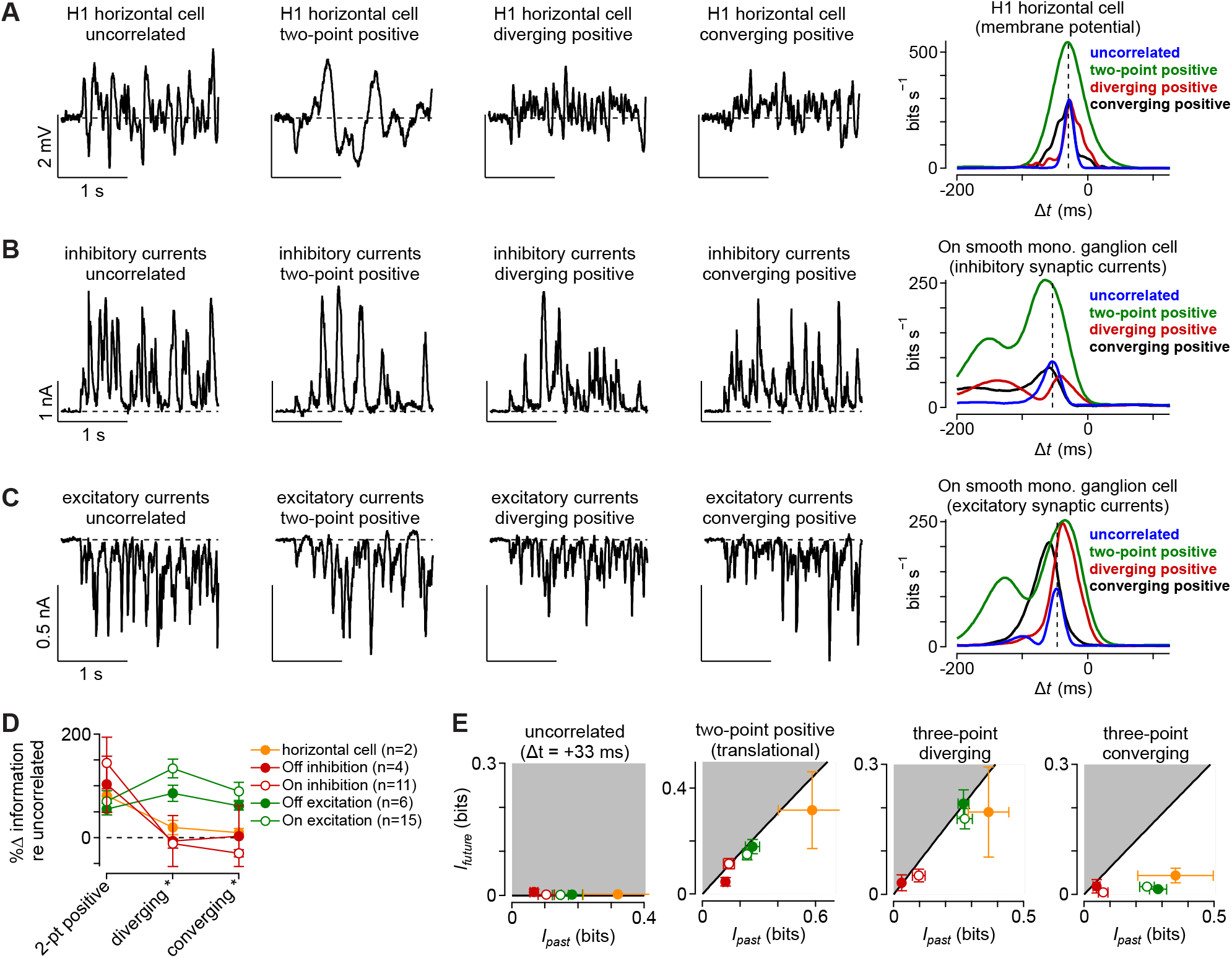
Information encoding varies with retinal processing stage. (A) Membrane potential in an H1 horizontal cell for four of the stimulus classes (*left*) and mutual information for the stimuli in the same cell (*right*). Peak information was similar between the uncorrelated condition and the three-point motion conditions, but was much higher for the two-point condition. (B) Same as (A) for the inhibitory synaptic currents in an On smooth monostratified ganglion cell. These currents showed a similar pattern observed in the horizontal cell, with a high information rate for the two-point motion correlations and smaller rates for the other stimulus classes. (C) Excitatory synaptic currents for the same On smooth monostratified cell as in (B). Here, the pattern shifted, with three-point motion correlations producing higher information rates relative to the uncorrelated condition and with predictive shifts toward positive time lags for the two-point and diverging motion stimuli. (D) Mutual information for two-point, diverging, and converging motion correlations relative to the information for the uncorrelated stimulus. Diverging and converging correlations data are shown stimuli aligned with a cell’s preferred contrast—positive contrasts for On-type cells and negative contrasts for Off-type cells. (E) Encoding of predictive information (*I_future_*; *y*-axis) versus past information (*I_past_*; *x*-axis) for the four stimulus classes. The black solid line indicates the theoretical boundary for predictive information and the shaded region defines theoretically unattainable values. Circles and error bars indicate mean ± SEM.

Horizontal cells receive excitatory synaptic inputs directly from cone photoreceptors and thus provide a window into neural processing very early in the visual stream. We recorded responses from H1 horizontal cells to our stimulus set and computed the time-shifted mutual information between each cell’s membrane potential and the stimulus (see Methods). Pair-wise motion correlations produced much higher mutual information values than the uncorrelated stimulus (Figure 6A, D). The higher-order correlations, however, showed similar information values to the uncorrelated control condition, indicating that much of the sensitivity to these correlations emerged down-stream of the horizontal cells in the retinal circuit. A similar pattern was observed in the inhibitory synaptic input from amacrine cells to parasol and smooth monostratified ganglion cells (Figure 6B).

The excitatory synaptic input from diffuse bipolar cells to these ganglion cells showed a pattern that was distinct relative to the horizontal cells or amacrine cell inhibition. Peak information rates for two-point and higher-order motion correlations were significantly higher than the uncorrelated condition. Further, the two-point and diverging motion stimuli produced a large shift in the information rates towards future time lags just as was observed in the ganglion cell spike outputs. These results indicate that the sensitivity to higher-order motion correlations arises after the horizontal cells in the visual processing stream, emerging at some point between the synaptic inputs and synaptic outputs of diffuse bipolar cells.

We also estimated the efficiency with which these circuit components encoded future information using the information bottleneck paradigm (Figure 6E). Indeed, the pattern of information in the excitatory synaptic inputs mirrored that observed in the spike outputs of parasol and smooth monostratified ganglion cells—encoding of future information approached the optimal boundary for both pairwise motion and diverging motion correlations. Inhibitory synaptic input also approached the boundary for these stimuli, but the overall magnitude of encoded future information was lower (see Discussion). Further, future information encoding in H1 horizontal cells fell away from the boundary, indicating that this nearly ideal encoding of future information emerges primarily in the diffuse bipolar cell circuitry. We next investigated the contribution of neural mechanisms in diffuse bipolar cells to the prioritized encoding of information about the future motion trajectory.

### Circuit nonlinearities favor encoding of predictive information

Our synaptic recordings revealed diffuse bipolar cells as the principal source at which selectivity for future information encoding emerges in the primate retina. Two known mechanisms in diffuse bipolar cells could contribute to encoding predictive motion information—electrical coupling and nonlinear input-output functions (Kuo et al., 2016; Manookin et al., 2018; Appleby and Manookin, 2020; Turner and Rieke, 2016). Together these components could work to enhance encoding of future information by first increasing cellular responsiveness to correlated stimuli through electrical coupling and, second, discarding weaker responses from uncorrelated stimuli with the output nonlinearity.

We tested this hypothesis using a computational model of the bipolar cell subunits based on direct excitatory synaptic input recordings from parasol and smooth monostratified ganglion cells (Manookin et al., 2018; Appleby and Manookin, 2020). In this model, the shape of the output nonlinearity was modified by shifting the threshold at which an input begins to produce subunit responses (Figure 7; see Methods). Model outputs were estimated to pairwise motion and to three-point diverging and converging motion stimuli (contrast, 0.5). Past and future information were calculated for the model across a range of thresholds. To determine whether electrical coupling between bipolar cells contributed to encoding future information, we compared the subunit model to a model that was identical except that model sub-units shared a portion of their responses through coupling (*coupled subunit model*).

**Figure 7.**
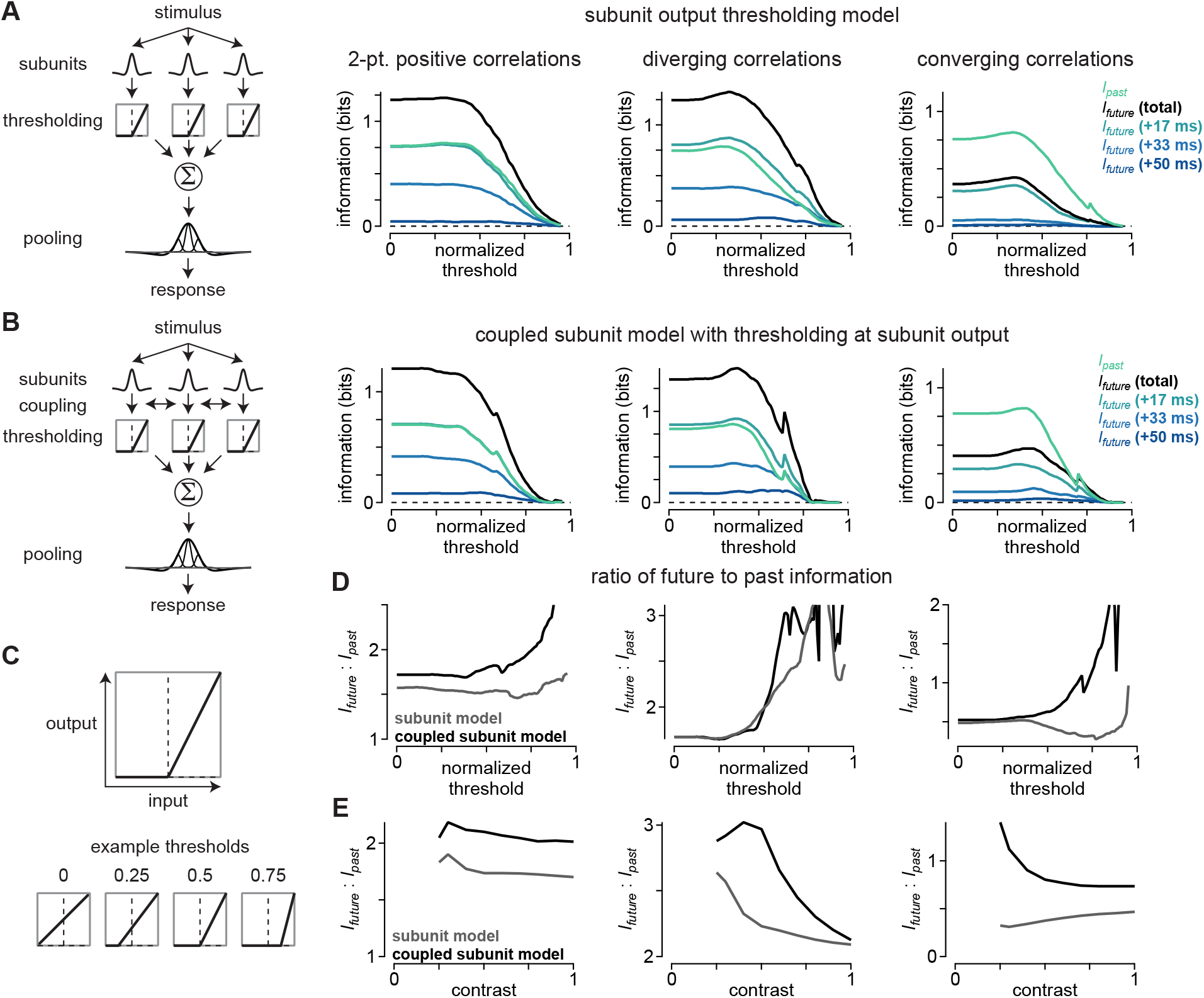
Subunit coupling and high output thresholds favor encoding of predictive information. (A) Model in which an output threshold was placed at the output of the model bipolar cell subunits. *Left*, The stimulus (contrast, 0.5) was filtered by receptive-field subunits. Subunit outputs were passed through an output threshold and then pooled at the level of the model ganglion cell at the ganglion cell output to determine circuit response. *Right*, Encoded information as a function of response threshold. Curves are shown for past information, total future information, and future information at three time lags. (B) The coupled subunit model was identical to the model in (A) except that model subunits were coupled and shared a portion of their responses prior to the output threshold. (C) Shapes of subunit input-output relationships at four different threshold levels. (D) Ratio of total future information to past information (*I_future_* : *I_past_*) for the two models as a function of threshold. Ratio of future to past information as a function of stimulus contrast for the two models at a threshold of 0.6. The coupled subunit model showed higher ratios at all contrasts.

We calculated the ratio of encoded future to past information to determine whether the output threshold or coupling would bias encoding of future information. The coupled subunit model showed a higher proportion of future versus past information encoded for all of the stimuli when the output threshold was in the range of ∼0.5–0.75. Thus, subunit coupling biased encoding of future over past information, but only when combined with a moderately high output threshold (Figure 7D).

To determine whether this pattern was consistent across a range of stimulus contrasts, we calculated the ratio of future to past information encoding across a range of contrasts (threshold, 0.6). Indeed, the coupled subunit model outperformed the model lacking coupling at all contrasts tested, indicating that coupling would enhance predictive motion encoding across a broad range of stimulus conditions (Figure 7E).

These modeling results support the hypothesis that known properties of diffuse bipolar cells—electrical coupling and input-output nonlinearities—shape the type of information that these cells encode. In summary, the correlations inherent in visual motion cause sequential stimulation of neighboring bipolar cells. As the motion excites a bipolar cell, a portion of the current from that cell passes to its neighbors. This shared current means that the neighboring cells will depolarize more as the motion stimulus passes through their receptive fields than if coupling were absent. Similar potentiation from electrical coupling will not occur for stimuli lacking spatiotemporal correlations, as sequential stimulation of neighboring neurons is required for this mechanism to be effective. Response thresholds contribute by discarding weak responses, which, due to the effects of network coupling, tend to lack strong spatiotemporal correlations. Thus, these mechanisms bias transmission of correlated inputs which, in the case of motion stimuli, contain information about the likely future trajectory of a moving object. Furthermore, our modeling results extend previous work on these neural mechanisms to demonstrate that they bias neural networks to both pairwise and higher-order motion correlations (Kuo et al., 2016; Manookin et al., 2018).

## DISCUSSION

The processing of light stimuli by the visual system involves unavoidable delays. These delays are particularly problematic for neural circuits tasked with encoding motion in the environment. Here, we studied four ganglion cell pathways in the primate retina that compensate for these delays by utilizing spatiotemporal correlations to predict future motion (Figure 3). Analysis of the types of encoded information revealed that these cells represented nearly all of the available predictive information for two-point and diverging motion in their spike outputs (Figure 4). Below, we summarize some of the principal implications of these findings.

### Efficient versus predictive encoding

Our study addresses how early visual circuits utilize correlations in incoming stimuli to predict motion. In the literature, ‘prediction’ is used in at least two different ways that are, in many ways, diametrically opposed to each other. In the context of efficient neural coding, prediction is a tool for compressing incoming sensory signals—neurons discard information that is predictable from past inputs and principally encode only those features that vary from that prediction. Thus, the encoding of stimulus features depends inversely on their likelihood of occuring in the environment (Brenner et al., 2000; Fairhall et al., 2001; Hosoya et al., 2005; Sharpee et al., 2006; Rao and Ballard, 1999; Johnston et al., 2019).

The second definition of predictive coding, consistent with the usage in the present study, indicates that stimulus information should be encoded based on its salience to animal behavior (Bialek et al., 2006; Palmer et al., 2015; Chalk et al., 2018; Salisbury and Palmer, 2016). In this paradigm, the information that can be used to estimate future states of the environment carries the greatest importance for guiding animal behavior and, thus, should be prioritized during neural encoding—this information is also termed predictive information.

Distinguishing between these hypotheses can be difficult as both paradigms can make similar predictions for empirical data. For example, the results of our experiment to test encoding at different stimulus contrasts can be interpreted to support both paradigms (Figure 5). At low contrast, the ratio of encoded future-to-past information was greater than at high contrast. Classical efficient coding predicts that as the signal-to-noise ratio increases, as with increasing contrast, the circuit should increasingly decorrelate the input. This decorrelation would remove some of the correlations in the stimulus needed to predict the future motion trajectory (Atick and Redlich, 1992; van Hateren, 1992; Atick and Redlich, 1990; Srinivasan et al., 1982; Dan et al., 1996; Fairhall et al., 2001; Sharpee et al., 2006; Brenner et al., 2000). Thus, the relative amount of future information would decrease at high contrast.

These data can also be interpreted through the lens of the framework in which information about the future is prioritized. Future information is prioritized and thus encoding of this information approaches the physical limit at low contrast. At high contrast, the circuit more readily encodes information about the past and the ratio of future-to-past information decreases (Bialek et al., 2006; Palmer et al., 2015; Chalk et al., 2018; Salisbury and Palmer, 2016). While both paradigms can explain these results, we favor the latter interpretation in which future information is prioritized during encoding, as this paradigm lends itself to straightforward interpretation in the context of the electrical coupling and nonlinear input-output functions of the diffuse bipolar cell networks—the origin of predictive motion encoding the primate retina.

Information that can be used to predict the future position of a moving object contains stronger spatiotemporal correlations than past information that is non-predictive. Electrical coupling and nonlinear synaptic transmission in the bipolar cell network produces stronger responses to correlated versus uncorrelated stimuli (Kuo et al., 2016; Manookin et al., 2018; Appleby and Manookin, 2020). Thus due to electrical coupling, at low contrast the more strongly correlated future information will drive the bipolar cell network harder than non-predictive information about the past that has weaker correlations. The weaker uncorrelated stimuli will then be discarded by the synaptic output nonlinearity whereas the stronger predictive information will be transmitted. Furthermore, stimuli with weaker correlations (e.g., non-predictive past information) will have a better chance of making it through the output nonlinearity at higher contrasts, which would explain the decrease in the ratio of future-to-past information at high contrast.

However, efficient coding and predictive coding are not mutually exclusive hypotheses. Neural circuits can remove predictable correlations in sensory inputs (i.e., efficient coding) while still preserving information needed to predict the trajectory of a moving object (i.e., predictive coding). Indeed, theoretical and empirical evidence indicates that removal of redundant information and encoding of information about the future are both critical components of neural coding in dynamic environments (Chalk et al., 2018; Hosoya et al., 2005).

A classic example of efficient coding is the difference-of-Gaussians receptive-field structure in the early visual system, which discards the low spatial frequencies that are common in nature—removing this predictable structure through decorrelation allows visual circuits to encode surprising features that vary from the prediction (Atick and Redlich, 1990; Srinivasan et al., 1982). However, this decorrelation of the spatial frequency information does not eliminate the phase information in the input image. This is a critical distinction because spatial frequency information is a statistical measure of an ensemble of natural images and, as such, is biased toward the average image. The phase information, however, is specific to individual images and can be used to identify features in those images. Thus, neural mechanisms could utilize this phase information in estimating future states of the environment (i.e., predictive coding) while, simultaneously, removing redundant spatial frequency information through decorrelation (i.e., efficient coding).

### Inhibitory synaptic inputs show weak information rates for higher-order correlations

Our inhibitory synaptic recordings showed relatively modest time-shifted mutual information values for higher-order motion correlations, yet amacrine cells receive synaptic input directly from bipolar cells, and bipolar cell output showed sensitivity to these correlations (Figure 6). This apparent discrepancy likely arose from the fact that parasol and smooth monostratified cells primarily receive inhibitory synaptic inputs from amacrine cells with the opposite contrast polarity. For example, an Off-type amacrine cell provides the dominant inhibitory synaptic inputs to On-type parasol ganglion cells (Cafaro and Rieke, 2010, 2013). However, for our synaptic input recordings, we made the decision to only record higher-order motion stimuli that aligned with the preferred contrast polarity of the recorded ganglion cells. This was done to ensure that we could obtain the large amount of data required for our analysis from each of the needed stimulus classes while also maintaining the stability of our whole-cell recordings.

We expect diverging and converging correlations that are opposite in polarity to the preferred contrast of the inhibitory amacrine cell will poorly drive predictive encoding. For example, triplet correlations that are positive contrasts would hyperpolarize Off-type bipolar cells such that the correlations contained in the stimulus would not exceed the threshold for synaptic release and would, thus, be discarded. Instead, these Off bipolar cells would depolarize most strongly at switches from positive to negative contrasts, similar to the response pattern expected for the uncorrelated stimuli. Such a response pattern would contain little information about the past motion trajectory of a positive-contrast stimulus. Indeed, this same pattern of relatively weak encoded information was observed in the ganglion cells to diverging and converging correlations that were of non-preferred contrast polarity (Figure 1, Figure 3). Thus, we would predict that inhibitory inputs to stimuli aligned with the preferred contrast polarity of these amacrine cells would show a similar pattern to our excitatory synaptic input recordings.

### Sensitivity to higher-order correlations arises early in the primate visual stream

The asymmetrical distribution of bright and dark intensities in nature produce correlations between three or more points in space and time during visual motion (Field, 1987; Dong and Atick, 1995; Fitzgerald et al., 2011; Hu and Victor, 2010; Ratliff et al., 2010). Indeed, humans and other animals utilize these higher-order correlations in estimating motion (Nitzany et al., 2017; Clark et al., 2014). Sensitivity to these correlations arise early in the visual stream in fruit flies—at the level of tangential cells in the lobula plate, if not earlier (Leonhardt et al., 2016). In humans and non-human primates, this sensitivity was thought to arise first in the primary visual cortex (Nitzany et al., 2017; Clark et al., 2014). Our findings indicate, however, that sensitivity to higher-order spatiotemporal correlations arises much earlier than previously thought— it is present in the synaptic outputs of diffuse bipolar cells at the second synapse of the visual stream. Thus, it appears that both flies and primates, with visual systems that evolved independently, employ a similar strategy of extracting and encoding these correlations early in visual processing (Clark et al., 2014; Leonhardt et al., 2016).

The common structure of natural scenes suggests that many sighted animals are faced with the problem of estimating motion from both first-order and higher-order correlations. While the results presented here indicate that at least four neural pathways in the retina contribute to this important function in primates, further work is needed to determine the generality of this neural computation across species and whether strategies for predicting motion from these correlations varies with ecological niche. Moreover, motion estimation is one of many prediction problems that animals must solve in order to survive and reproduce. The neural mechanisms supporting motion prediction identified in this work—subunit pooling and nonlinear thresholding—are features shared across vertebrate evolution in the retina and other brain regions. Thus, we hope that our work will guide future studies aimed at understanding the neural mechanisms that solve these diverse computational problems.

## ACKNOWLEDGEMENTS

We thank Shellee Cunnington for technical assistance. Tissue was provided by the Tissue Distribution Program at the Washington National Primate Research Center (WaNPRC; supported through NIH grant P51 OD-010425), and we thank the WaNPRC staff, particularly Chris English and Audrey Baldessari, for making these experiments possible. Chris Chen assisted in tissue preparation. We thank Siwei Wang and Gabrielle Gutierrez for helpful discussions, and Chuan-Chin Chiao for supporting B.L. and A.H. This work was supported in part by grants from the NIH (NEI R01-EY027323 to M.B.M.; NEI R01-EY029247 to E.J.C., F.R., and M.B.M.; NEI R01-EY028542 to F.R.; NEI P30-EY001730 to the Vision Core), Research to Prevent Blindness Unrestricted Grant (to the University of Washington Department of Ophthalmology), Latham Vision Research Innovation Award (to M.B.M.), the Alcon Young Investigator Award (to M.B.M.), the Taiwanese Ministry of Science and Technology (108-2813-C-007-085-B to A.H.), and a travel award to B.L. and A.H. from the Taiwanese Ministry of Education (Chuan-Chin Chiao, PI).

## AUTHOR CONTRIBUTIONS

M.B.M. designed the study. F.R. and M.B.M. performed the experiments. B.L., A.H., F.R., and M.B.M. designed the analytical techniques. M.B.M. wrote the analysis code. B.L., A.H., F.R., and M.B.M. interpreted the results and wrote the paper.

## METHODS

These experiments were done using an *in vitro* preparation of the macaque monkey retina from three different macaque species of either sex (Macaca *fascicularis*, *mulatta*, and *nemestrina*). Tissues were obtained from terminally anesthetized animals that were made available through the Tissue Distribution Program of the National Primate Research Center at the University of Washington. All procedures were approved by the University of Washington Institutional Animal Care and Use Committee.

Cellular recordings were performed in a whole-mount retinal preparation. To aid with cellular visualization, horizontal cell recordings were made with pigment-epithelium removed, and all other recordings were done with the pigment-epithelium intact.

### Visual stimuli

Visual stimuli were generated using the Stage software package (http://stage-vss.github.io) and displayed on a customized digital light projector designed specifically for studying visual processing in old-world primates (Appleby and Manookin, 2019, 2020). All stimuli in this work were updated at 60 Hz. Stimuli were presented at medium to high photopic light levels with average L/M-cone photoisomerization rates (R*) of ∼1.5 × 10^4^ – 5.0 × 10^5^ s^−1^. Contrast values are given in Michaelson contrast. "Glider" stimuli with distinct spatiotemporal correlations were generated as described in Jonathan Victor’s previous work (Hu and Victor, 2010; Nitzany et al., 2016).

### Mutual information calculations

Mutual information quantifies how much knowing the response (*R*) of a neuron can reduce the uncertainty about a presented stimulus (*S*). Neural responses were binned at the stimulus presentation rate (update rate, 60 Hz; bin width, 16.7 ms) for all analyses with the exception of the data in Figure 2, which were binned at 1 kHz (bin width, 1 ms). We estimated the amount of information that cellular responses at a particular time (*r_t_*) provided about the stimulus at time, *t′* (*s_t_′*), where *t′* = *t* + Δ*t* using Equation 1.

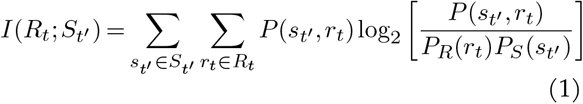

where *P_R_*(*r*) is the distribution of responses in a single cell, *P_S_* (*s*) is the stimulus distribution, and *P* (*s_t′_, r_t_*) is the joint distribution of stimuli presented at time *t′* and responses *r* observed at time *t*. In other words, responses were fixed in time, the stimulus was shifted for each time bin between ±0.5 seconds, and the mutual information was computed at each of these time shifts (Palmer et al., 2015; Shannon, 1948; Rieke et al., 1997; Bialek, 2012; Chien, 2017; Chen et al., 2017). The time-shifted mutual information analysis assumes that neighboring time bins can be treated independently. This may not be the case due to serial correlations in the stimulus and neural responses, but a full treatment of these correlations would require a prohibitive amount of data.

Response levels were determined by either counting the number of spikes occurring in a time bin (extracellular recordings) or distributing analog responses across six levels (current-clamp and voltage-clamp recordings). Varying the number of response levels between 4–8 for the analog data did not noticeably affect the information computations, so we used a value near the center of that range. These mutual information calculations required converting the spatial dimensions of our stimuli into a single value for each time bin. We did this by first identifying the four spatial regions of the stimulus that were centered over the receptive field. Each of the 16 possible stimulus patterns for those four regions was assigned a value between 0–15.

To correct for inflation error in our mutual information calculations, stimulus and response data were randomly subsampled at fractions of 0.5–1.0. Information was calculated 50 times for each fraction and information was extrapolated for an infinite number of samples (Strong et al., 1998; Palmer et al., 2015). Inflation error was estimated as the standard deviation of the information calculated for data subsampled at a fraction of 0.5 divided by 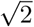. An error value below was a criterion for inclusion in our study and, in all cases, error values were <6 × 10^−3^ bits spike^−1^ (Palmer et al., 2015).

### Information bottleneck calculations

We used the information bottleneck method to determine the optimal encoding/compression schemes for the stimuli used in this study (Tishby et al., 1999). The idea is that the past stimulus, *S_past_*, can be compressed into a representation (*Z*) that retains sufficient information that can be used to estimate future motion trajectories (*S_future_*). The optimal mapping between *S_past_* and its compressed representation *Z* was determined by solving Equation 2:

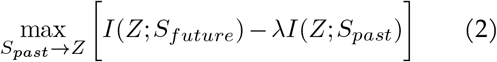

where the Lagrangian multiplier, *λ*, determines how much information is retained about the past stimulus. Varying the value of *λ* reflects different tradeoffs between representing the greatest possible amount of information about the future and producing a more compressed representation. Large values of *λ* exemplify the former case and, thus, poor representations of future information are strongly penalized. For small *λ* values, the goal is to achieve compression and we are, thus, more likely to ignore disparities in how *Z* represents future information (Bialek, 2012).

Given the relatively low dimensionality of our stimuli (16 possible states on a single frame), solving Equation 2 for each of the possible compression schemes was a computationally tractable problem. Thus, we computed *I*(*Z*; *S_future_*) and *I*(*Z*; *S_past_*) for every possible stimulus mapping in *Z*. This process was repeated for three future time lags of +17, +33, and +50 ms. Detailed methods for computing the bottleneck are given elsewhere (Slonim and Tishby, 2000; Tishby et al., 1999). Briefly, the algorithm was initialized with *Z* ≡ *S_past_* and *Z* was modified by merging different combinations of the 16 possible stimulus states. Mutual information *I*(*Z*; *S_future_*) and *I*(*Z*; *S_past_*) was calculated from the corresponding joint probability matrices, *P* (*s_future_, z*) and *P* (*s_past_, z*) for each of the possible mergings using Equation 1.

### Computing the temporal lag in visual processing

We calculated the spatiotemporal receptive fields of our cells to estimate the temporal lag between the presentation of light stimuli at the level of the cone photoreceptors and the output of the retinal neurons in our study. Receptive fields were determined from the average stimulus preceding a spike (Chichilnisky, 2001):

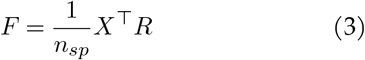

where *X* is a matrix with the rows containing time samples and the columns containing the spatiotemporal sequence in the previous 0.5 seconds. *R* is a column vector containing the spike counts in the corresponding time bins of *X* and *n_sp_* is the spike count. The time lag was then defined as the time bin at which the spatiotemporal filter (*F*) reached a maximum (On-type cells) or minimum (Off-type cells).

### Stimulus autocorrelation calculations

To confirm that the three-point motion stimuli lacked net two-point spatiotemporal correlations, we calculated the autocorrelation functions of these stimuli. Stimulus autocorrelation for a stimulus sequence, *X*, was determined by first subtracting the mean stimulus (*μ*) and then taking the outer product of the result with itself.

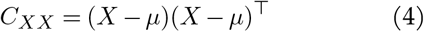

where *X* is a matrix in which the rows are time points and the columns are the spatiotemporal sequence of contrasts occurring in the preceding second.

### Sensitivity index calculations

We evaluated a cell’s ability to accurately detect a change in information using the sensitivity index (*d′*), which measures the amount of overlap between the information distributions in the original spike train and after shifting spike timing:

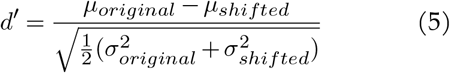

where *μ_original_* and *μ_shifted_* are the mean of the original and shifted information distributions and 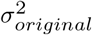 and 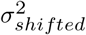 are the variances of those distributions, respectively.

### Quantification and statistical analysis

All statistical analyses were performed in MATLAB (R2018b, Mathworks). Final figures were created in MATLAB (version R2018b), Igor Pro (version 8), and Adobe Illustrator.

## SUPPLEMENTARY MATERIALS

### Time-shifted information calculations

**Figure S1.**
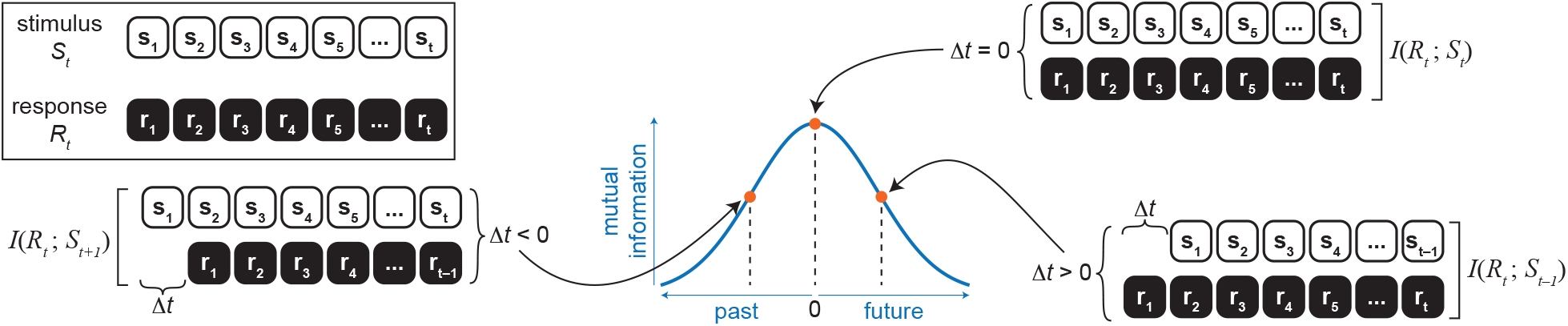
Overview of time-shifted mutual information calculations. The stimulus sequence was shifted relative to the response and mutual information was computed between the stimulus and response. This was done for time shifts between ±0.5 seconds.

### Information calculations for local motion correlations

**Figure S2.**
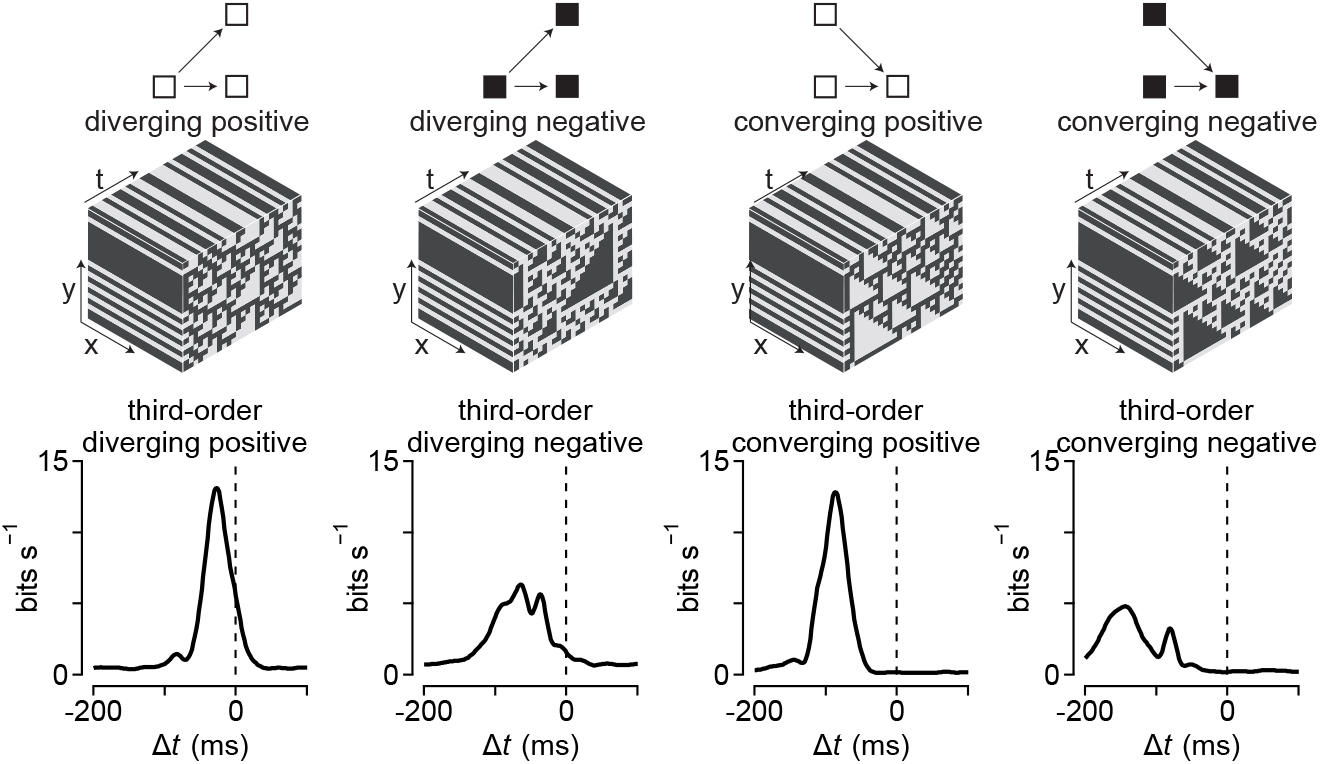
Local motion signals show temporally shifted profiles. Third-order spatiotemporal correlations were extracted directly from stimulus sequences. The time-shifted mutual information profile for diverging correlations showed shifts toward future time points, whereas converging information was shifted toward past time lags.

We performed an alternative analysis to determine the time-shifted mutual information in our cells. For this analysis, the stimulus was collapsed to a single value in each time bin by directly computing the motion correlations as described previously (Nitzany and Victor, 2014). Briefly, the strength of spatiotemporal correlations in each time bin *C*(*t*) was determined from the product of contrasts at points in space and time that corresponded to the space and time lags in the motion stimulus and subtracting the spatially reversed contrast values as described in Equation 6.

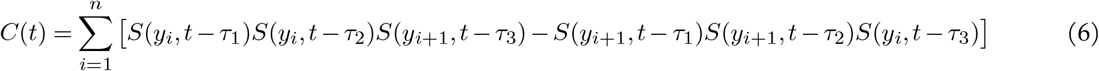

This approach did not provide a means to compare information in the motion stimuli to the uncorrelated control stimulus, so we did not use it as our primary information measure. However, the technique did serve as another demonstration that the cells under study show predictive encoding for triplet diverging motion correlations.

